# Preventing neutrophil from oxygen exposure allows their basal state maintenance

**DOI:** 10.1101/2020.10.15.340919

**Authors:** Louise Injarabian, Quentin Giai Gianetto, Véronique Witko-Sarsat, Benoit S Marteyn

## Abstract

Neutrophils are the most abundant circulating white blood cells and are central players of the innate immune response. During their lifecycle, neutrophils mainly evolve under low oxygen conditions (0.1 - 4% O_2_) to which they are well adapted. Neutrophils are atypical cells since they are mainly glycolytic, and highly susceptible to oxygen-exposure, which induces their activation and death, through mechanisms which remain currently elusive. Nevertheless, nearly all studies conducted on neutrophils are carried out under atmospheric oxygen (21%), corresponding to hyperoxic conditions. Here we investigated the impact of hyperoxia during neutrophil purification and culture on neutrophil viability, activation and cytosolic protein content. Neutrophil hyper-activation (CD62L shedding) is induced during culture under hyperoxic conditions (24h), compared to neutrophils cultured under anoxic conditions. In addition, we show that maintaining neutrophils in autologous plasma is the most suitable strategy to maintain their basal state.

Our results show that manipulating neutrophils under hyperoxic conditions leads to the loss of ~100 cytosolic proteins during purification, while it does not lead to an immediate impact on neutrophils activation (CD11b^high^, CD54^high^, CD62L^low^) or viability (DAPI^+^). We identified two clusters of proteins belonging to the cholesterol metabolism and to the complement and coagulation cascade pathways, which are highly susceptible to neutrophil oxygen-exposure during their purification.

In conclusion, preserving neutrophil from oxygen-exposure during their manipulation – purification and culture- is recommended to avoid their experimental activation and for preserving a large set of cytosolic proteins from alteration.

## Introduction

Polymorphonuclear neutrophils (neutrophils, PMNs) are the most abundant leukocytes in the blood circulation (2.5-7.5×10^9^ PMNs/L). Neutrophils are atypical and fully differentiated cells, with a polylobal nucleus (3-5 lobes), a granule-rich cytosol, a low mitochondrial content and produce most of their energy using glycolysis (1). Neutrophils differentiate from hematopoietic stem cells (HSCs, CD34+) in the bone marrow, in an hypoxic compartment (1.3 - 4.1% O_2_) (2). Under basal conditions, 10% of the mature bone marrow neutrophils are released into the blood plasma fraction, containing low amounts of dissolved oxygen (~ 0.1-1.2%) (3) (4). In conclusion, neutrophils are adapted to low concentrations of oxygen during their lifecycle, facing more elevated oxygen tensions only while transmigrating into perfused tissues, during inflammation or infection processes. Under basal conditions, tissues oxygen level is <10% O_2_ (reviewed in (5)), whereas hypoxia (<1% O_2_) is induced during inflammation and infection (inflammatory hypoxia and infectious hypoxia) (6) (7).

The impact of hypoxia on neutrophils has been well documented (reviewed in (8) (9)). Moreover, our group and others have demonstrated that increased neutrophil viability is maintained under anoxic conditions (0% O_2_) (10) (11), although in these studies neutrophils were transiently exposed to atmospheric oxygen during their purification prior their culture under controlled-oxygen conditions. Conversely, molecular mechanisms mediating the deleterious impact of hyperoxia on neutrophil physiology remain unclear.

In this report, we investigated the impact of hyperoxia (21%) during neutrophil purification and culture on their viability and activation, demonstrating the importance of preventing cells from oxygen-exposure for the maintenance of neutrophil basal state.

## Results and Discussion

### Differential oxygen availability during neutrophil purification induces changes in protein abundance

We previously demonstrated that when neutrophils were purified under hyperoxic conditions (21% O_2_, atmospheric air), neutrophil viability was maintained upon subsequent culture under anoxic conditions (11). We therefore hypothesized that avoiding any exposure to oxygen during the purification and manipulation of neutrophils may be beneficial to prevent cell death induction or cell activation. Thus, an innovative method was developed consisting in purifying neutrophils form peripheral human blood samples under oxygen-free conditions (anoxia).

Upon reception, human blood samples were immediately transferred into an anoxic chamber or kept under atmospheric conditions (Fig. 1A). Neutrophils were purified in parallel using the same conditions on a Percoll® gradient and, as a control, a rapid purification method was used (MACSxpress® Whole Blood Neutrophil Isolation Kit, Miltenyi) (Fig. 1A). We demonstrated by flow cytometry that the viability of neutrophils just after the purification was not significantly different between anoxic and hyperoxic purifications (Fig. 1B, *p*>0.05). However, the viability of neutrophil was slightly increased in both conditions, compared to the viability of neutrophils purified with a rapid method (Fig. 1B, *p*<0.001 (hyperoxia) and *p*<0.001 (anoxia)). We further assessed neutrophil activation upon anoxic or hyperoxic purifications and observed no significant difference using CD54, CD11b and CD62L cell-surface markers (Fig S1A-C, *p*>0.05). It has to be noticed here that the purification of neutrophils on a Percoll® gradient slightly tend to activate neutrophils (CD62L shedding) in both conditions, compared to the rapid purification method (Fig. S1C, *p*<0.01).

**Figure 1.**
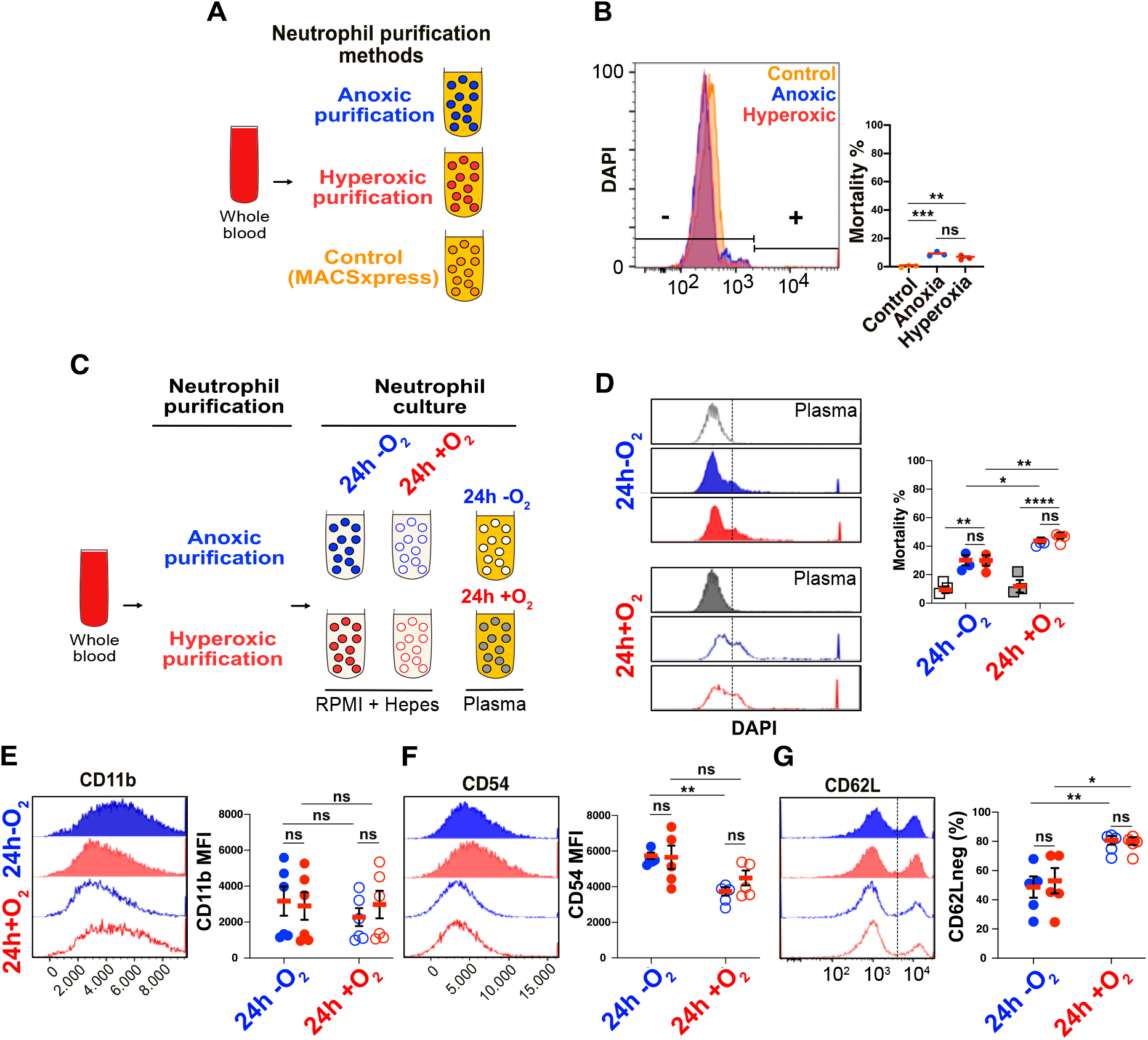
Respective impact of neutrophil purification and culture in anoxia on cell viability maintenance and activation limitation. (**A**) Experimental plan description: comparison of neutrophils purified in anoxia (blue), hyperoxia (red) or with a rapid-purification method MACSxpress (control; orange). (**B**) Flow cytometry assessment of neutrophils mortality (DAPI^+^ cells, (**p=0.0037; ***p=0.0003)). (**C**) Experimental plan description: comparison of neutrophils purified in anoxia or hyperoxia, exposed to anoxia or hyperoxia for 24h in RPMI or in autologous plasma (control). (**D**). Assessment of DAPI^+^ neutrophils after anoxic or hyperoxia purification and incubation (24h) in RPMI and autologous plasma measured with flow cytometry. **p=0.001/0.002; ***p<0.0001; *p=0.0263; **p=0.0049). Assessment of (**E**) CD11b^high^ (**F**) CD54^high^;**p=0.0072), and (**G**) CD62L^low^ (**p=0.0035, *p=0.0114) neutrophils after anoxic or hyperoxia purification and incubation (24h) in RPMI measured with flow cytometry.

### Combining neutrophil purification and culture under anoxic conditions maintains their viability and limits their activation

We previously demonstrated that the plasma fraction of the blood contains a limited amount of dissolved oxygen (<1.2% O_2_ (4)). Accordingly, we set up a new method consisting in limiting oxygen-exposure during neutrophil culture and compared its impact on the phenotype of neutrophils purified in anoxia or hyperoxia (Fig. 1C). This strategy allowed to evaluate the respective impact of oxygen-exposure during neutrophil purification and culture on their viability and activation. Neutrophils are short-lived cells, with an estimated life expectancy ranging from few hours to few days (12) (13). To observe changes in neutrophil viability and activation, anoxia/hyperoxia-purified neutrophils were cultured in anoxia or hyperoxia for 24h in a previously optimized culture medium (11). As a control, anoxia-purified neutrophils were cultured in autologous plasma in anoxia/hyperoxia (Fig. 1C).

First, we demonstrated that neutrophil survival was increased when cultured in plasma, irrespective of their exposition to oxygen (<10% mortality, Fig. 1D and Fig. S3), suggesting that in addition to oxygen-exposure prevention, other plasma components protect neutrophils from cell death. Based on this observation, further investigations will be required to identify these beneficial components.

When neutrophils were cultured in RMPI+Hepes medium, we confirmed that their mortality was increased in hyperoxia, as observed previously (11), regardless of the purification conditions (anoxia/hyperoxia) (Fig. 1D, *p*<0.01 and *p*<0.01 (Fig. S3)). The corollary of this result is that no significant impact of the purification conditions (anoxia/hyperoxia) on neutrophil survival was observed, regardless of the culture conditions (*p*>0.05, Fig. 1D and Fig. S3). In conclusion, culturing neutrophil under anoxic conditions is the most important parameter to maintain their viability. Above all, maintaining neutrophil in plasma is essential to maintain their quiescence and viability.

We subsequently assessed neutrophil activation in all tested conditions. No significant difference was observed using the CD11b marker, probably due to the large variability of the results (Fig. 1E, *p*>0.05). Conversely, we demonstrated that combining neutrophil purification and culture under anoxia allows to significantly reduce the cell-surface exposure of Icam-1 (CD54) (Fig. 1F, *p*<0.01). Culturing under anoxia neutrophils purified under hyperoxia do not lead to such a significant protection, as compared to hyperoxic cultures (Fig. 1F, *p*>0.05). Last, we demonstrated that neutrophil culture under anoxia significantly limits CD62L shedding, regardless of the purification conditions (anoxia/hyperoxia) (Fig. 1G, *p*<0.05 and *p*<0.01 respectively), which is in accordance with previous studies on neutrophils from mice and rabbits (14) (15) (16).

In conclusion, we have demonstrated here that neutrophil culture under anoxia is crucial to maintain their viability and limit their activation *in vitro* (Fig. 1D-G). We have shown that purifying neutrophils under anoxia do not lead to significant changes regarding these two parameters (Fig. 1D-G). Beyond neutrophil cell-surface marker abundance and viability assessment, we further investigated the impact of the purification method on neutrophil cytosolic protein content.

### Neutrophil purification under hyperoxic conditions (21% O_2_) leads to the loss of ~100 cytosolic proteins

We hypothesized that the stability of neutrophil cytosolic oxidation-sensitive proteins may be affected by a transient exposure to oxygen (hyperoxic conditions) of neutrophils during their purification. This hypothesis was validated using a proteomic analysis, revealing that the neutrophil cytosolic protein content was largely modified upon their purification under anoxic or hyperoxic conditions. Mass-spectrometry sample quality control using principal component analysis (PCA) indicated very little variation between samples enabling the study of the majority of cytosolic proteins (Fig. 2A-B, n=3). Indeed, most of the cytosolic proteins identified by mass spectrometry were present in all replicates (Fig. 2A, 1741 proteins upon anoxic purification and 1940 proteins upon hyperoxic purification). Strikingly, a large set of proteins (148 proteins) was more abundant in the cytosol of neutrophils purified under anoxia, while 56 proteins were significantly more abundant in the cytosol of neutrophils purified under hyperoxia (Fig. 2C and Fig. S4 and S5). More importantly, only 4 proteins were uniquely present in the cytosol of neutrophils purified under hyperoxic conditions (Fig. S4), while 57 proteins were uniquely present in the cytosol of neutrophils purified under anoxic conditions (Fig. S5). We performed a pathway enrichment analysis of uniquely abundant proteins using Search Tool for the Retrieval of Interacting Genes/Proteins (STRING) database (11.0). Interestingly, STRING analysis did not reveal any metabolic pathway specifically expressed in neutrophils purified under hyperoxic conditions (Fig. 2D and S4), while the cholesterol pathway (KEGG ID : hsa04979 composed of APOA1, APOA2, APOA4 and APOC3) and the complement activation and coagulation cascades (KEGG ID: hsa04610 composed of C3, CFH, SERPINF2 and VTN) were uniquely enriched when neutrophils were purified under anoxic conditions (Fig. 2E and S5).

**Figure 2.**
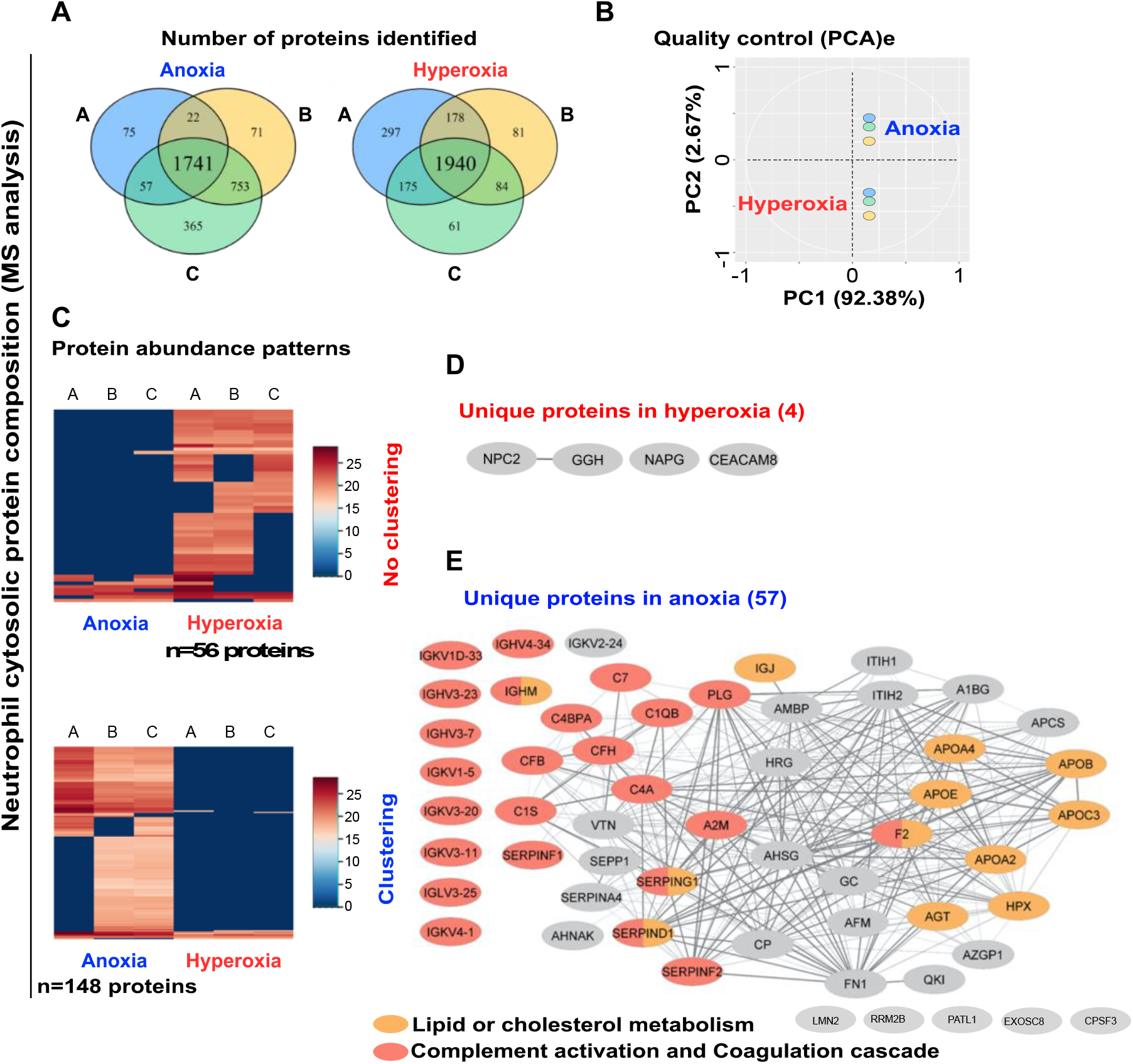
Neutrophil cytosolic protein composition is altered during their purification in hyperoxia, compared to anoxia. (**A**) Commonly identified proteins in all three samples (A, B, C) in anoxia and hyperoxia. (**B**) Data quality control assessment using principal component analysis (PCA) of cytosolic protein samples of neutrophils purified in anoxia or hyperoxia after mass-spectrometry analysis. (**C**) Heatmap representing differential protein abundance between anoxia- and hyperoxia-purified human neutrophils. (**D**) Pathway enrichment analysis of uniquely identified proteins in hyperoxic conditions using Database for Annotation, Visualization, and Integrated Discovery (DAVID) and Cytoscape for network visualization. (**E**) Enrichment of the lipid and cholesterol metabolism (GO0043691, GO0001523, GO0042157, GO0008203) and the complement and coagulation cascade (GO0006958/hsa04610) pathways identified with DAVID and visualized with Cytoscape in anoxia-purified neutrophils. Enrichment p-values for the GO terms and hsa pathways are p(GO0043691; Reverse cholesterol transport)=9.10×10^−6^, p(GO0001523; Retinoid metabolic process)=1.77×10^−5^, p(GO0042157; Lipoprotein metabolic processes)=2.6×10^−4^, p(GO0008203; Cholesterol metabolic process)=3.76×10^−4^, p(GO0006958; Complement activation, classic pathway)=3.87×10^−22^, p(hsa04610; Complement and coagulation cascade)=5.57×10^−12^.

In conclusion, preserving neutrophils from hyperoxia during their purification, manipulation and physiological study is a critical parameter to maintain cells under a basal state. Our results confirmed the beneficial effect of the neutrophil purification and culture under anoxic conditions. New metabolic and signaling pathways have been revealed in this study, which open new doors in the study of neutrophil physiology.

## Acknowledgments

The authors wish to thank Morgane Le Gall from the Institut Cochin Proteomic platfrom for her assistance in data mining. This work was supported by the ANR JCJC grant (ANR-17-CE15-0012) (B.S.M.).

## Notes

### Competing Interest Statement

The authors have declared no competing interest.

