## Supplementary Figures and Methods for "Preventing neutrophil from oxygen exposure allows their basal state maintenance"

### Injarabian *et al*, Supplementary informations

#### 1- Supplementary Figures

**Supplementary Figure 1.** Complementary results to Fig. 1A-B. Flow cytometry assessment of neutrophils cell-surface marker adundance :(**A**) CD54$^{high}$**, (B**) CD11b$^{high}$ and (**C**) CD62L$^{low}$ (***p*=0.0011, ***p*=0.0037) after anoxic, hyperoxic or MACSxpress (control) purification.

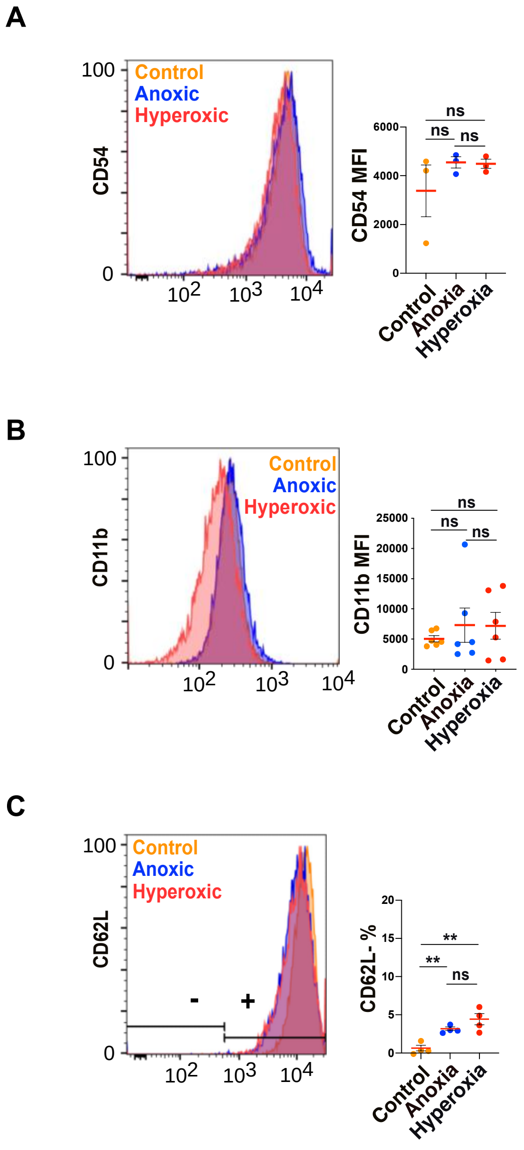

**Supplementary Figure 2.** ANOVA analysis of flow cytometry results obtained on neutrophils purified under anoxic and hyperoxic conditions and incubated in anoxia or hyperoxia for 24h in RPMI1640 (complemented with 10mM Hepes and 10% autologous plasma) or 100% autologous plasma (control) as described in Fig. 2B-E. The proportion of DAPI$^{+}$ cells (**A**), the MFI of CD54 (**B**), CD11b (**C**) and the percentage of CD62L$^{low}$ cells (**D**) between differently purified (anoxic or hyperoxic) neutrophils were compared and significant differences indicated.

**A**

| Tukey's multiple comparisons test | Predicted (LS) mean diff. | 95.00% CI of diff. | Significant? | Summary | Adjusted P Value |
|---|---|---|---|---|---|
| 24h-O2 | | | | | |
| Plasma vs. Anoxic purification | -23.87 | -43.90 to -3.845 | Yes | * | 0.0179 |
| Plasma vs. Hyperoxic purification | -25.73 | -45.76 to -5.705 | Yes | * | 0.0107 |
| Anoxic purification vs. Hyperoxic purification | -1.86 | -19.20 to 15.48 | No | ns | 0.9603 |
| 24h+O2 | | | | | |
| Plasma vs. Anoxic purification | -43.31 | -63.34 to -23.29 | Yes | **** | <0.0001 |
| Plasma vs. Hyperoxic purification | -44.35 | -64.38 to -24.33 | Yes | **** | <0.0001 |
| Anoxic purification vs. Hyperoxic purification | -1.04 | -18.38 to 16.30 | No | ns | 0.9874 |

**B**

| Tukey's multiple comparisons test | Predicted (LS) mean diff. | 95.00% CI of diff. | Significant? | Summary | Adjusted P Value |
|---|---|---|---|---|---|
| 24h-O2 | | | | | |
| Anoxic purif. vs. Hyperoxic purif. | 83.4 | -2005 to 2172 | No | ns | 0.9946 |
| 24h+O2 | | | | | |
| Anoxic purif. vs. Hyperoxic purif. | -752 | -2841 to 1337 | No | ns | 0.6484 |

**C**

| Tukey's multiple comparisons test | Mean Diff. | 95.00% CI of diff. | Significant? | Summary | Adjusted P Value |
|---|---|---|---|---|---|
| 24h-O2 | | | | | |
| Anoxic purif. vs. Hyperoxic purif. | 267.2 | -2056 to 2590 | No | ns | 0.9567 |
| 24h+O2 | | | | | |
| Anoxic purif. vs. Hyperoxic purif. | -704.7 | -3027 to 1618 | No | ns | 0.7372 |

**D**

| Tukey's multiple comparisons test | Mean Diff. | 95.00% CI of diff. | Significant? | Summary | Adjusted P Value |
|---|---|---|---|---|---|
| 24h-O2 | | | | | |
| Anoxic purif. vs. Hyperoxic purif. | -4.32 | -27.00 to 18.36 | No | ns | 0.8806 |
| 24h+O2 | | | | | |
| Anoxic purif. vs. Hyperoxic purif. | 0.46 | -22.22 to 23.14 | No | ns | 0.9985 |

**Supplementary Figure 3.** ANOVA analysis of flow cytometry results obtained on neutrophils purified under anoxic and hyperoxic conditions and incubated in anoxia or hyperoxia for 24h in RPMI1640 (complemented with 10mM Hepes and 10% autologous plasma) or 100% autologous plasma (control) as described in Fig. 2B-E. The proportion of DAPI$^+$ cells (**A**), the MFI of CD54 (**B**), CD11b (**C**) and the percentage of CD62L$^{low}$ cells (**D**) between differently incubated (in anoxia or hyperoxia) neutrophils were compared and significant differences indicated.

**A**

| Sidak's multiple comparisons test | Mean Diff. | 95.00% CI of diff. | Significant? | Summary | Adjusted P Value |
|---|---|---|---|---|---|
| 24h-O2 - 24h+O2 | | | | | |
| Plasma | -2.487 | -14.47 to 9.499 | No | ns | 0.9238 |
| Anoxic purification | -13.5 | -25.49 to -1.514 | Yes | * | 0.0263 |
| Normoxic purification | -17.47 | -29.45 to -5.481 | Yes | ** | 0.0049 |

**B**

| Sidak's multiple comparisons test | Mean Diff. | 95.00% CI of diff. | Significant? | Summary | Adjusted P Value |
|---|---|---|---|---|---|
| 24h-O2 - 24h+O2 | | | | | |
| Anoxic purification | 2003 | 553.4 to 3452 | Yes | ** | 0.0072 |
| Normoxic purification | 1167 | -282.0 to 2616 | No | ns | 0.1246 |

**C**

| Sidak's multiple comparisons test | Mean Diff. | 95.00% CI of diff. | Significant? | Summary | Adjusted P Value |
|---|---|---|---|---|---|
| 24h-O2 - 24h+O2 | | | | | |
| Anoxic purification | -31.98 | -53.01 to -10.95 | Yes | ** | 0.0035 |
| Normoxic purification | -27.2 | -48.23 to -6.166 | Yes | * | 0.0114 |

**Supplementary Figure 4.** Uniquely identified proteins in hyperoxia-purified neutrophils (4 proteins). The number of unique and specific peptides identifies in each replicate (A, B, C) of neutrophils purified under anoxic or hyperoxic conditions. We excluded proteins that had a unique peptide value >0 in one of the replicates in anoxia. Table contains the Uniprot number, Protein code, Protein name, Entrez Gene ID and the number of unique peptides found in each sample

| | | | | | Anoxia | | | Hyperoxia | | |
|---|---|---|---|---|---|---|---|---|---|---|
| **Uniprot** | **Protein code** | **Protein name** | **Entrez Gene ID** | **Gene name** | **A** | **B** | **C** | **A** | **B** | **C** |
| J3KMY5 | J3KMY5_HUMAN | Epididymal secretory protein E1 | 10577 | NPC2 | 0 | 0 | 0 | 3 | 4 | 2 |
| A0A087X2E2 | A0A087X2E2_HUMAN | deleted entry | 1088 | CEACAM8 | 0 | 0 | 0 | 1 | 0 | 3 |
| Q6FHY4 | Q6FHY4_HUMAN | N-ethylmaleimide-sensitive factor attachment protein, gamma, isoform CRA_b | 8774 | NAPG | 0 | 0 | 0 | 4 | 2 | 1 |
| A8K335 | A8K335_HUMAN | Folate gamma-glutamyl hydrolase | 8836 | GGH | 0 | 0 | 0 | 3 | 3 | 1 |

**Supplementary Figure 5.** Uniquely identified proteins in anoxia-purified neutrophils (57 proteins). The number of unique and specific peptides identifies in each replicate (A, B, C) of neutrophils purified under anoxic or hyperoxic conditions. We excluded proteins that had a unique peptide value >0 in one of the replicates in hyperoxia. Table contains the Uniprot number, Protein code, Protein name, Entrez Gene ID and the number of unique peptides found in each sample. Proteins belonging to the "Lipid or cholesterol metabolism" and the "Complement and Coagulation cascade" described in the Fig. 1F-G are highlighted in pink and orange, respectively. If proteins participate in both pathways then ½ is in orange and ½ in pink. Non-clustering proteins (5) are separated in the bottom of the table.

| | | | | | **Anoxia** | | | **Hyperoxia** | | |
|---|---|---|---|---|---|---|---|---|---|---|
| **Uniprot** | **Protein code** | **Protein name** | **Entrez Gene ID** | **Gene name** | **A** | **B** | **C** | **A** | **B** | **C** |
| P01023 | A2MG_HUMAN | Alpha-2-macroglobulin | 2 | A2M | 60 | 46 | 42 | 0 | 0 | 0 |
| A0A0G2JPR0 | A0A0G2JPR0_HUMAN | C4a anaphylatoxin | 720 | C4A | 47 | 14 | 19 | 0 | 0 | 0 |
| A0A087X2C0 | A0A087X2C0_HUMAN | immunoglobulin heavy constant mu(IGHM) | 3507 | IGHM | 17 | 16 | 14 | 0 | 0 | 0 |
| V9GYM3 | APOA2/V9GYM3_HUMAN | Apolipoprotein A-II | 336 | APOA2 | 7 | 5 | 5 | 0 | 0 | 0 |
| P06727 | APOA4_HUMAN | Apolipoprotein A-IV | 337 | APOA4 | 23 | 11 | 16 | 0 | 0 | 0 |
| D6RF35 | D6RF35_HUMAN | Gc-globulin | 2638 | GC | 18 | 10 | 6 | 0 | 0 | 0 |
| P02790 | HEMO_HUMAN | Hemopexin | 3263 | HPX | 17 | 9 | 9 | 0 | 0 | 0 |
| A8K5A4 | A8K5A4_HUMAN | cDNA FLJ76826, highly similar to Homo sapiens ceruloplasmin (ferroxidase) (CP), mRNA | 1356 | CP | 27 | 10 | 14 | 0 | 0 | 0 |
| B4E1Z4 | B4E1Z4_HUMAN | C3/C5 convertase | 629 | CFB | 27 | 3 | 10 | 0 | 0 | 0 |
| Q5T985 | Q5T985_HUMAN | Inter-alpha-trypsin inhibitor heavy chain H2 | 3698 | ITIH2 | 20 | 7 | 14 | 0 | 0 | 0 |
| B7ZLE5 | B7ZLE5_HUMAN | Fibronectin | 2335 | FN1 | 39 | 16 | 35 | 0 | 0 | 0 |
| A0A0F7T737 | A0A0F7T737_HUMAN | IGHV4-34 protein | 28395 | IGHV4-34 | 3 | 2 | 1 | 0 | 0 | 0 |
| B2R8I2 | B2R8I2_HUMAN | cDNA, FLJ93914, highly similar to Homo sapiens histidine-rich glycoprotein (HRG), mRNA | 3273 | HRG | 10 | 13 | 10 | 0 | 0 | 0 |
| Q5UGI6 | Q5UGI6_HUMAN | Serine/cysteine proteinase inhibitor clade G member 1 splice variant 2 | 710 | SERPING1 | 13 | 5 | 5 | 0 | 0 | 0 |

| | | | | | | | | | | |
|---|---|---|---|---|---|---|---|---|---|---|
| B7Z549 | B7Z549_HUMAN | cDNA FLJ56821, highly similar to Inter-alpha-trypsin inhibitor heavy chain H1 | 3697 | ITIH1 | 17 | 3 | 8 | 0 | 0 | 0 |
| A0A125QYY5 | A0A125QYY5_HUMAN | GCT-A9 light chain variable region | 28896 | IGKV1D-33 | 2 | 2 | 2 | 0 | 0 | 0 |
| P00747 | PLMN_HUMAN | Plasminogen | 5340 | PLG | 17 | 9 | 8 | 0 | 0 | 0 |
| Q0ZCH9 | Q0ZCH9_HUMAN | Immunglobulin heavy chain variable region | 28452 | IGHV3-7 | 4 | 2 | 2 | 0 | 0 | 0 |
| C0JYY2 | C0JYY2_HUMAN | Apolipoprotein B (Including Ag(X) antigen) | 338 | APOB | 66 | 34 | 95 | 0 | 0 | 0 |
| Q9UL78 | Q9UL78_HUMAN | Myosin-reactive immunoglobulin light chain variable region | 28912 | IGKV3-20 | 4 | 4 | 4 | 0 | 0 | 0 |
| A0N5G1 | A0N5G1_HUMAN | Rheumatoid factor C6 light chain | 28299 | IGKV1-5 | 2 | 2 | 2 | 0 | 0 | 0 |
| A3KPE2 | APOC3/A3KPE2_HUMAN | Apolipoprotein C-III | 345 | APOC3 | 3 | 2 | 2 | 0 | 0 | 0 |
| P00734 | THRB_HUMAN | Prothrombin | 2147 | F2 | 11 | 2 | 2 | 0 | 0 | 0 |
| C9JV77 | C9JV77_HUMAN | Alpha-2-HS-glycoprotein | 197 | AHSG | 6 | 4 | 3 | 0 | 0 | 0 |
| P43652 | AFAM_HUMAN | Afamin | 173 | AFM | 13 | 1 | 3 | 0 | 0 | 0 |
| A0A0C4DH68 | A0A0C4DH68_HUMAN | Immunoglobulin kappa variable 2-24 | 28923 | IGKV2-24 | 2 | 1 | 2 | 0 | 0 | 0 |
| V9HWD8 | V9HWD8_HUMAN | Epididymis secretory sperm binding protein Li 163pA | 1 | A1BG | 10 | 6 | 3 | 0 | 0 | 0 |
| A0A140VK00 | A0A140VK00_HUMAN | Testicular tissue protein Li 227 | 563 | AZGP1 | 12 | 3 | 5 | 0 | 0 | 0 |
| Q09666 | AHNK_HUMAN | Neuroblast differentiation-associated protein AHNAK | 79026 | AHNAK | 3 | 1 | 78 | 0 | 0 | 0 |
| P05546 | HEP2_HUMAN | Heparin cofactor 2 | 3053 | SERPIND1 | 14 | 4 | 6 | 0 | 0 | 0 |
| D6RHJ6 | D6RHJ6_HUMAN | Immunoglobulin J chain | 3512 | IGJ | 4 | 2 | 2 | 0 | 0 | 0 |
| B4E1B3 | B4E1B3_HUMAN | Angiotensin 1-10 | 183 | AGT | 8 | 4 | 5 | 0 | 0 | 0 |
| A0A024R962 | CFH/A0A024R962_HUMAN | Complement Factor F/HCG40889, isoform CRA_b | 3075 | CFH | 17 | 4 | 9 | 0 | 0 | 0 |
| A2KBC8 | A2KBC8_HUMAN | Anti-TeTox scFv | 28442 | IGHV3-23 | 6 | 4 | 5 | 0 | 0 | 0 |
| P08697 | SERPINF2/A2AP_HUMAN | Serpin Family F Member 2/Alpha-2-antiplasmin | 5345 | SERPINF2 | 8 | 4 | 4 | 0 | 0 | 0 |
| V9HWP0 | V9HWP0_HUMAN | Pentaxin | 325 | APCS | 4 | 1 | 2 | 0 | 0 | 0 |
| B4E1D8 | B4E1D8_HUMAN | cDNA FLJ51597, highly similar to C4b-binding protein alpha chain | 722 | C4BPA | 9 | 1 | 3 | 0 | 0 | 0 |
| A0A0X9TD47 | A0A0X9TD47_HUMAN | MS-D1 light chain variable region | 28914 | IGKV3-11 | 2 | 2 | 2 | 0 | 0 | 0 |
| A0A0S2Z3D5 | A0A0S2Z3D5_HUMAN | Apolipoprotein E isoform 1 | 348 | APOE | 9 | 1 | 4 | 0 | 0 | 0 |
| P02760 | AMBP_HUMAN | Protein AMBP | 259 | AMBP | 4 | 1 | 4 | 0 | 0 | 0 |
| D9ZGG2 | VTN/D9ZGG2_HUMAN | Vitronectin | 7448 | VTN | 6 | 3 | 4 | 0 | 0 | 0 |

| | | | | | | | | | | |
|---|---|---|---|---|---|---|---|---|---|---|
| A0A087X232 | A0A087X232_HUMAN | Complement C1s subcomponent | 716 | C1S | 10 | 2 | 2 | 0 | 0 | 0 |
| Q5NV90 | Q5NV90_HUMAN | V2-17 protein | 28793 | IGLV3-25 | 3 | 2 | 2 | 0 | 0 | 0 |
| A0A140VKF3 | A0A140VKF3_HUMAN | Testis tissue sperm-binding protein Li 70n | 5176 | SERPINF1 | 11 | 3 | 0 | 0 | 0 | 0 |
| P01625 | KV402_HUMAN | Immunoglobulin kappa variable 4-1 | 28908 | IGKV4-1 | 3 | 1 | 2 | 0 | 0 | 0 |
| B2R6W1 | B2R6W1_HUMAN | cDNA, FLJ93143, highly similar to Homo sapiens complement component 7 (C7), mRNA | 730 | C7 | 8 | 1 | 4 | 0 | 0 | 0 |
| A0A0A0MSV6 | A0A0A0MSV6_HUMAN | Complement C1q subcomponent subunit B | 713 | C1QB | 3 | 0 | 2 | 0 | 0 | 0 |
| B2R815 | B2R815_HUMAN | cDNA, FLJ93695, highly similar to Homo sapiens serpin peptidase inhibitor, clade A (alpha-1 antiproteinase, antitrypsin), member 4 (SERPINA4), mRNA | 5267 | SERPINA4 | 7 | 0 | 2 | 0 | 0 | 0 |
| P49908 | SEPP1_HUMAN | Selenoprotein P | 6414 | SEPP1 | 2 | 1 | 2 | 0 | 0 | 0 |
| H0YGD6 | H0YGD6_HUMAN | Protein quaking | 9444 | QKI | 0 | 2 | 3 | 0 | 0 | 0 |
| Q96ST3 | SIN3A_HUMAN | Paired amphipathic helix protein Sin3a | 25942 | SIN3A | 0 | 4 | 7 | 0 | 0 | 0 |
| K7EKR9 | K7EKR9_HUMAN | Ribonucleoprotein PTB-binding 1 | 125950 | RAVER1 | 0 | 3 | 3 | 0 | 0 | 0 |
| H0YAV1 | H0YAV1_HUMAN | Ribonucleoside-diphosphate reductase | 50484 | RRM2B | 0 | 3 | 3 | 0 | 0 | 0 |
| B4DQR2 | B4DQR2_HUMAN | cDNA FLJ57562, highly similar to Cleavage and polyadenylation specificityfactor 73 kDa subunit | 51692 | CPSF3 | 0 | 2 | 3 | 0 | 0 | 0 |
| Q86TB9-2 | PATL1_HUMAN | Protein PAT1 homolog 1 | 219988 | PATL1 | 0 | 2 | 3 | 0 | 0 | 0 |
| J9JID7 | J9JID7_HUMAN | deleted entry | 84823 | LMN2 | 4 | 5 | 1 | 0 | 0 | 0 |
| Q96B26 | EXOS8_HUMAN | Exosome complex component RRP43 | 11340 | EXOSC8 | 0 | 2 | 2 | 0 | 0 | 0 |

#### 2- Supplementary Extended Methods

*Neutrophil purification*

All participants gave written, informed consent in accordance with the Declaration of Helsinki principles. Peripheral human blood was collected from healthy patients at the ICAReB service of the Pasteur Institute (authorization DC No.2008-68) and from the Etablissement Français du Sang (EFS) de Strasbourg (authorization n°ALC/PIL/DIR/AJR/FO/606). Human blood samples were collected from the antecubital vein into tubes or blood collection bags containing sodium citrate (3,8% final) as an anticoagulant.

For percoll-gradient human neutrophil purification, whole blood samples were centrifuged at 1800 rpm for 20 minutes without a break. Platelet rich plasma (PRP) was collected and centrifuged at 3800 rpm for 20 min to form platelet poor plasma (PPP). Blood cells were resuspended in NaCl 0.9% and dextran sulfate (0.72% final). After 30 min sedimentation, the leukocyte-containing upper layer was centrifuged at 300 x *g* for 10 min. The resuspended pellet was separated on a 42% Percoll-plasma (GE Healthcare) gradient by centrifugation at 800 x *g* for 20 min. Neutrophils were collected from the pellet with remaining red blood, which was removed using CD235a (glycophorin) microbeads (negative selection, Miltenyi Biotec). Anoxic purification steps were performed in the anoxic chamber using oxygen-free media.

For MACSxpress purification of human neutrophils we used MACSxpress Whole Blood Neutrophil Isolation Kit, human (Miltenyi Biotec), following the manufacturer procedure. Remaining red blood cells were removed using CD235a (glycophorin) microbeads.

*Neutrophil culture*

Purified neutrophils were centrifuged 10 min 300 x *g* and resuspended at $1.10^6$ cells/ml in RPMI1640 medium (Gibco) supplemented with 10mM Hepes buffer (ThermoFisher) and 10% autologous plasma. Neutrophils were plated into a 24-well plate at 1ml/well in in the presence or absence of oxygen (21% and 0% $O_2$).

*Flow cytometry*

Cell viability and activation was measured by flow cytometry. Cells in culture were resuspended and centrifuged 10 min at 300 x *g*. Cell-containing pellets were subsequently resuspended in 300 μL PBS + 2 mM EDTA and incubated for 15 min at RT in the presence of the following fluorescent markers (1/100 dilution), as indicated: 4′,6-diamidino-2-phenylindole (DAPI), CD11b-PE; CD54-APC, CD62L-FITC. Labelled cells were analyzed with an 8-color cell analyzer BD FACSCanto™ (BD Biosciences). Data mining was achieved with the FlowJo software (FlowJo, LLC). Data were further analyzed using Prism 8.0 software (GraphPad) for statistical analyses.

*Mass spectrometry*

*Cell extracts preparation*. Human neutrophils purified under hyperoxic or anoxic conditions were pelleted and resuspended in ice-cold PBS at $27.10^6$ cells/ml containing Diisopropylfluorophosphate (DFP, Sigma-Aldrich) (0.5 µL/mL) and incubated on ice for 15 min. The cells were centrifuged at 300 x *g* for 10 min at 4°C. Pellets were resuspended at $20.10^6$ cells/mL in relaxation buffer 1x containing 3 mM PMSF, 1 mM orthovanadate, 400 µM pepstatin, 400 µM leupeptin, 1 mM ATP, 1 mM EDTA. Resuspended cells were introduced into a cavitation bomb chamber. Pressure was stabilized at 350 psi for 20 minutes with nitrogen. Cell membranes were disrupted by a slow gas expansion. Cell lysates were collected in tubes containing 1.25 mM EGTA (1) (2).

Neutrophil cytosolic fractions were isolated by centrifugation. First, nuclei and remaining cells were removed by centrifugating samples at 500 x *g* for 15 min at 4°C. Then, neutrophil granules and organites were removed by centrifugating samples at 37000 x *g* for 1.5h at 4°C (Beckmann, 11x32 MM PC tubes ref 343778, TLA102 rotor, TL100 ultracentrifuge or equivalent).

Protein concentration was determined in each sample and 5 µg of protein was run on a 12%SDS-PAGE to select homogeneous samples. Proteins (45µg) were digested for 14 hours at 37°C with 1 µg trypsin (Promega), and were fractionated by strong cationic exchange (SCX) StageTips. Mass spectrometry analyses were performed on a U3000 RSLC nano-LC-system coupled to an LTQ Orbitrap-Velos mass spectrometer (Thermo Fisher Scientific). The data were analyzed using MaxQuant version 1.5.2.8. The database used was a concatenation of human sequences from the Uniprot–Swissprot database (Uniprot, release 2015-02) and a list of contaminant sequences from MaxQuant. The identification false discovery rate (FDR) was kept below 1% on both peptides and proteins. Quantification of each identified protein was performed by using the MaxLFQ algorithm available in MaxQuant (4). Label-free protein quantification (LFQ) was carried out using both unique and razor peptides. At least two such peptides were required for LFQ quantification of a protein.

*Bioinformatic analysis of proteomics data*

Proteins identified in the reverse and contaminant databases and proteins “only identified by site” (with an identification score too low - not exceeding the 1% FDR threshold) were first discarded. Then, proteins exhibiting fewer than 2 LFQ intensities in at least one condition (either Anoxia or Hyperoxia) were discarded from the list to ensure a minimum of replicability in quantified values.

After log2 transformation of the LFQ intensities of the leftover proteins, Principal Component Analysis (PCA) was performed from the proteins without missing values in all the samples (856 proteins). Logged LFQ intensities in each sample have been centred and scaled before analysis. The variables factor map (Figure 1) represents each sample projected on the first and second component of the PCA. It allows visualizing clusters of correlated samples. Of note, the LFQ intensities of samples are globally strongly correlated whatever the condition, such that the first principal component explains most of the variance (92%), while the second component distinguishes Anoxic samples from Hyperoxic samples (2.6% of explained variance). This shows the good reproducibility of the quantified values.

The differential analysis highlighting more (148 proteins) or less (56 proteins) abundant proteins in Anoxia than in Hyperoxia was carried out using an in-house R analysis pipeline based on the R packages DAPAR (5), imp4p (6), limma (7) and cp4p (8). Statistical enrichment tests were performed from these proteins to highlight major Gene Ontology terms or KEGG pathways that may be affected using DAVID software v6.8 (9) by selecting the set of quantified proteins among all samples as background. The protein-protein interaction graph was determined from STRING v11 (10) and visualized using Cytoscape v3.8.0 (11) with the plugins stringApp (12) and Omics Visualizer (13). Only “exclusive” proteins with unique peptides identified in a condition (Anoxia or Hyperoxia) and no peptide identified in the other condition, were represented. Widths of the edges correspond to the *combined score* of STRING reflecting the confidence placed in each interaction. The proteins highlighted as "Lipid or Cholesterol metabolism" and "Complement activation and coagulation cascades" were determined from enriched annotation clusters using the fuzzy heuristic clustering method of DAVID v6.8 (10).
